# Oxidative regulation of TDP-43 self-association by a β-to-α conformational switch

**DOI:** 10.1101/2023.08.29.555361

**Authors:** Jinge Gu, Xiaoming Zhou, Lillian Sutherland, Masato Kato, Klaudia Jaczynska, Josep Rizo, Steven L. McKnight

**Affiliations:** Department of Biochemistry, UT Southwestern Medical Center, 5323 Harry Hines Blvd., Dallas, Texas 75235; Department of Biophysics, UT Southwestern Medical Center, 5323 Harry Hines Blvd., Dallas, Texas 75235; Institute for Quantum Life Science, National Institutes for Quantum Science and Technology (QST), 4-9-1, Anagawa, Inage-ku, Chiba, JAPAN 263-8555

**Author notes:** corresponding author –. authors contributing equally (Jinge Gu, Xiaoming Zhou).

## Abstract

An evolutionarily conserved region of the TDP-43 low complexity domain twenty residues in length can adopt either an α-helical or β-strand conformation. When in the latter conformation, TDP-43 self-associates via the formation of a labile, cross-β structure. Self-association can be monitored via the formation of phase separated protein droplets. Exposure of droplets to hydrogen peroxide leads to oxidation of conserved methionine residues distributed throughout the low complexity domain. Oxidation disassembles the cross-β structure, thus eliminating both self-association and phase separation. Here we demonstrate that this process reciprocally enables formation of α-helical structure in precisely the same region formerly functioning to facilitate β-strand mediated self-association. We further observe that the α-helical conformation allows interaction with a lipid-like detergent, and that exposure to lipids enhances the β-to-α conformational switch. We hypothesize that regulation of this oxidative switch will prove to be important to the control of localized translation within vertebrate cells. The experimental observations reported herein were heavily reliant on studies of 1,6-hexanediol, a chemical agent that selectively dissolves labile structures formed via the self-association of protein domains of low sequence complexity. This aliphatic alcohol is shown to exert its dissociative activity primarily via hydrogen bonding interactions with carbonyl oxygen atoms of the polypeptide backbone. Such observations underscore the central importance of backbone-mediated protein:protein interactions that facilitate the self-association and phase separation of low complexity domains.

**Significance Statement:** The TDP-43 protein is a constituent of RNA granules involved in regulated translation. TDP-43 contains a C-terminal domain of 150 amino acids of low sequence complexity conspicuously decorated with ten methionine residues. An evolutionarily conserved region (ECR) of 20 residues within this domain can adopt either of two forms of labile secondary structure. Under normal conditions wherein methionine residues are reduced, the ECR forms a labile cross-β structure that enables RNA granule condensation. Upon methionine oxidation, the ECR undergoes a conformational switch to become an α-helix incompatible with self-association and granule integrity. Oxidation of the TDP-43 low complexity domain is hypothesized to occur proximal to mitochondria, thus facilitating dissolution of RNA granules and activation of localized translation.

## Introduction

The genomic coding regions of all life forms specify the formation of proteins that fold into stable, three-dimensional structures. The variety and complexity of folded proteins are enabled by polypeptide sequences that make use of most or all of life’s twenty amino acids. An unusual class of proteins is characterized by sequences containing no more than a small subset of amino acids. These protein domains of low sequence complexity are unable to fold into stable, three-dimensional structures. Upwards of 10-20% of eukaryotic proteomes are composed of these intrinsically disordered sequences.

The phenylalanine-glycine (FG)-rich proteins occupying the central channel of nuclear pores represent prominent examples of low complexity (LC) domains. Nearly two decades ago an inventive study gave evidence that a purified nuclear pore FG domain was able to become phase separated from aqueous solution in a gel-like state (1, 2). That this representation of the protein might be of biologic substance was favored by evidence that FG domain hydrogels allowed highly selective penetration of an importin β-like protein known to shuttle cargo through nuclear pores.

Conceptually and experimentally analogous observations on a transport pore specific to peroxisomal membranes have recently been reported (3). Similar findings have also been observed for the LC domains of numerous RNA-binding proteins and intermediate filament proteins (4-6). All of these varied domains of low sequence complexity are able to self-associate in a manner leading to phase separation, thus creating biochemical assays for the study of this enigmatic class of proteins.

The LC domains of nuclear pore proteins, RNA-binding proteins and intermediate filaments share the common feature of being enriched in aromatic amino acids. Mutational alteration of the phenylalanine residues of nuclear pore proteins, or the tyrosine residues of RNA-binding proteins, commonly impede both hydrogel formation and biological function (2, 5). The LC domains of two proteins relevant to neurodegenerative disease, ataxin-2 and TAR DNA-binding protein 43 (TDP-43), display patterns of amino acids distinctly different from prototypic LC domains. Both ataxin-2 and TDP-43 replace aromatic amino acids with methionine, leading to the discovery that methionine oxidation can reversibly inhibit phase separation (7-9). Variants of yeast ataxin-2, wherein methionine residues were replaced by either phenylalanine or tyrosine, are immune to oxidation-mediated inhibition of phase separation. Moreover, yeast cells expressing oxidation-resistant variants of the ataxin-2 protein are incapable of transducing a mitochondrion-derived signal normally employed to control activity of the target-of-rapamycin (TOR) complex and autophagy (8).

Efforts to distil an understanding of how the TDP-43 LC domain can reversibly phase separate as a function of methionine oxidation led to the discovery of a small region of 20 amino acids perfectly conserved in vertebrates for 500M years since the divergence of fish and humans (10). When incubated under physiologically normal conditions of monovalent salt and neutral pH, this region forms a labile, cross-β structure responsible for self-association and phase separation. Perplexingly, this same region alternatively adopts an α-helical structure when the TDP-43 LC domain is studied in the monomeric state under acidic conditions devoid of monovalent salt ions (11-13). Here we show that methionine oxidation triggers a β-to-α molecular switch that blocks self-association and simultaneously assists formation of a lipophilic, α-helical epitope. We hypothesize that this switch will prove to be of central importance to the control of localized translation in vertebrate cells.

## Results

### Aliphatic Alcohols Preferentially Melt Self-associated LC Domains

An early study contributed by the laboratory of Dr. Dirk Gorlich gave evidence that the permeability barrier of nuclear pores was compromised upon treatment of cultured HeLa cells with an aliphatic alcohol (14). Subsequent studies of two aliphatic alcohols sharing the identical chemical formula, 1,6-hexanediol (1,6-HD) and 2,5-hexanediol (2,5-HD), have been particularly useful to the study of protein domains of low sequence complexity. The value of these regioisomeric chemicals derives from the concordance of their activities on sub-cellular structures in living cells that are enriched with LC domains as compared with phase separated hydrogels and liquid-like droplets formed in test tubes from purified LC domains.

Despite sharing identical molecular formulas, these two chemicals display contrasting activities. When assayed in living cells, 1,6-HD melts the permeability barrier of nuclear pores (15), intermediate filaments (6, 16), and a wide variety of nuclear and cytoplasmic puncta not surrounded by investing membranes (6, 17-19). By contrast, 2,5-HD exhibits significantly diminished capacity to dissolve these cellular structures. The same differences in activity are observed *in vitro* when the two aliphatic alcohols are tested on either hydrogels or liquid-like droplets formed from purified, phase separated LC domains (6, 20).

How do these chemicals work, why is 1,6-HD more adept than 2,5-HD at reversing self-association of LC domains? To interrogate these questions, we initially compared the activities of the two aliphatic alcohols on two samples of phase separated proteins (Fig. 1 and *SI Appendix* Fig. S1). The first sample was liquid-like droplets formed from the LC domain of TDP-43. 1,6-HD melts these droplets at an EC_50_ concentration of 4%, which is considerably lower than the 8% half-maximal melting level observed for 2,5-HD (Fig. 1).

**Fig. 1.**
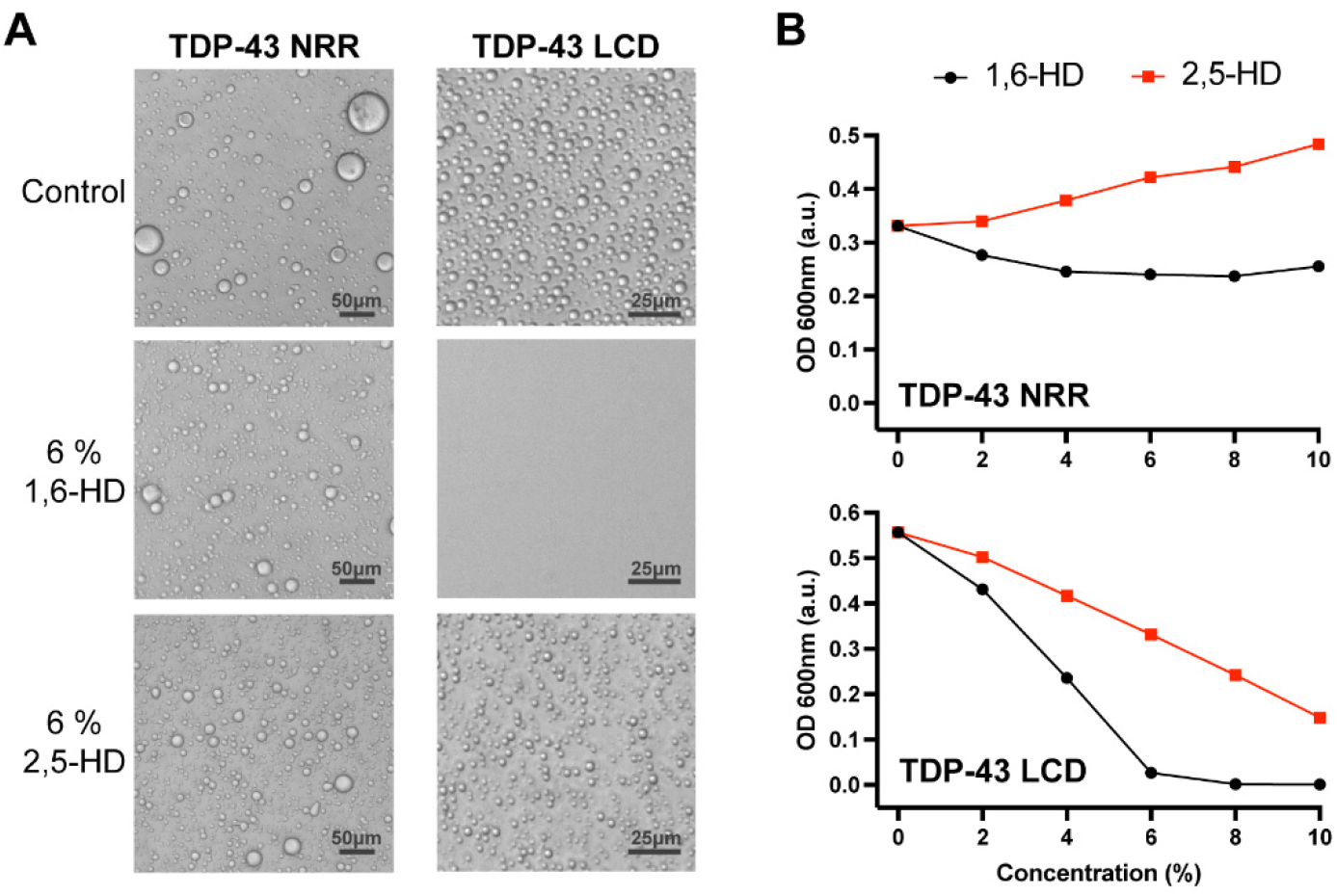
Effects of aliphatic alcohols on phase separated droplets formed from the structured and unstructured halves of the TDP-43 RNA-binding protein. (A) Protein fragments corresponding to residues 1-262 or 263-414 of the TDP-43 protein were expressed in bacteria, purified and incubated under conditions of neutral pH and physiologically normal monovalent salt ions. Both the N-terminal fragment (residues 1-262) bearing the structured N-terminal oligomerization domain and two RRM domains (NRR), and the C-terminal low complexity domain (LCD, residues 263-414) became phase separated in the form of spherical protein droplets under a protein concentration of 15 to 20 μM. 4% PEG-8,000 was added to the purified NRR to facilitate formation of liquid-like droplets. Such droplets were immune to the effects of both 1,6-hexanediol (1,6-HD) and the regioisomeric 2,5-hexanediol (2,5-HD) chemical. Droplets formed from the C-terminal LCD were fully melted upon exposure to 6% 1,6-HD, but left largely intact upon exposure to 6% 2,5-HD. (*B*) Phase separation by the TDP-43 NRR and LCD was quantified by measurements of turbidity. Neither aliphatic alcohol caused a reduction in turbidity of the sample formed by the TDP-43 NRR (upper panel). At all concentrations tested, 1,6-HD reduced turbidity more substantially than 2,5-HD for droplets formed from the TDP-43 LCD (lower panel).

The second sample was liquid-like droplets formed from a 262-residue segment of TDP-43 containing its N-terminal oligomerization domain and two RRM domains (21). All three of these domains represent well-folded segments of the TDP-43 protein. In this case droplet formation required addition of 4% polyethylene glycol (PEG) 8000. This form of protein phase separation is likely to be biologically irrelevant, yet PEG-assisted protein condensation is widely used by researchers in the phase separation field and represents a recommended method by field-leading scientists (22). Even upon exposure to 10% 1,6-HD, no evidence of droplet melting was observed for PEG8000-assisted condensates composed of the structured domains of TDP-43. These observations confirm published studies showing that 1,6-HD is incapable of melting liquid-like droplets formed upon incubation of the three structured domains of TDP-43 at high protein concentrations (21). Likewise, as shown in Figure 1B, 2,5-HD also failed to melt phase separated droplets formed by the structured regions of the TDP-43 protein (Fig. 1).

These observations help explain why 1,6-HD affects neither actin filaments nor microtubules, yet readily disassembles intermediate filaments (6). Actin filaments and microtubules are formed from exclusively structured proteins. By contrast, intermediate filament assembly is critically dependent upon LC domains that flank either end of the long, α-helical coiled-coil domains prototypic of intermediate filaments (IFs). Functional studies have confirmed that the LC head domains of IF proteins are required for filament assembly (23). As tested both *in vivo* and *in vitro*, 1,6-HD melts intermediate filaments more readily than its 2,5-HD regioisomer by inhibiting the cross-β self-association of the LC head domains (6, 16). This concept has recently been confirmed by resolution of the atomic structure of assembled vimentin intermediate filaments (24). The molecular structure of vimentin IFs reveals a central lumen formed by LC head domains organized in an amyloid-like, cross-β conformation.

### Aliphatic Alcohols Interact with Carbonyl Oxygen Atoms of the Polypeptide Backbone

We next employed NMR spectroscopy to probe protein interaction by the 1,6-HD and 2,5-HD aliphatic alcohols. Our aspiration in conducting these experiments was to discover differences in protein binding by the two chemicals. Since 1,6-HD dissolves phase separated droplets formed by the TDP-43 LC domain far more readily than 2,5-HD, any differences in protein binding by the two chemicals might yield useful mechanistic insight.

Protein samples corresponding to the LC domain of TDP-43, the well-folded SUMO1 protein, and the well-folded C2A and C2B domains of synaptotagmin1 were uniformly labeled with ^13^C and ^15^N, purified and subjected to various analyses by solution NMR spectroscopy. Protein binding by the two aliphatic alcohols was visualized in the form of ^13^C chemical shift changes of phenylalanine and tyrosine aromatic rings, aliphatic side chains including methionine, leucine, isoleucine and valine, and carbonyl groups of the polypeptide backbone (Fig. 2).

**Fig. 2.**
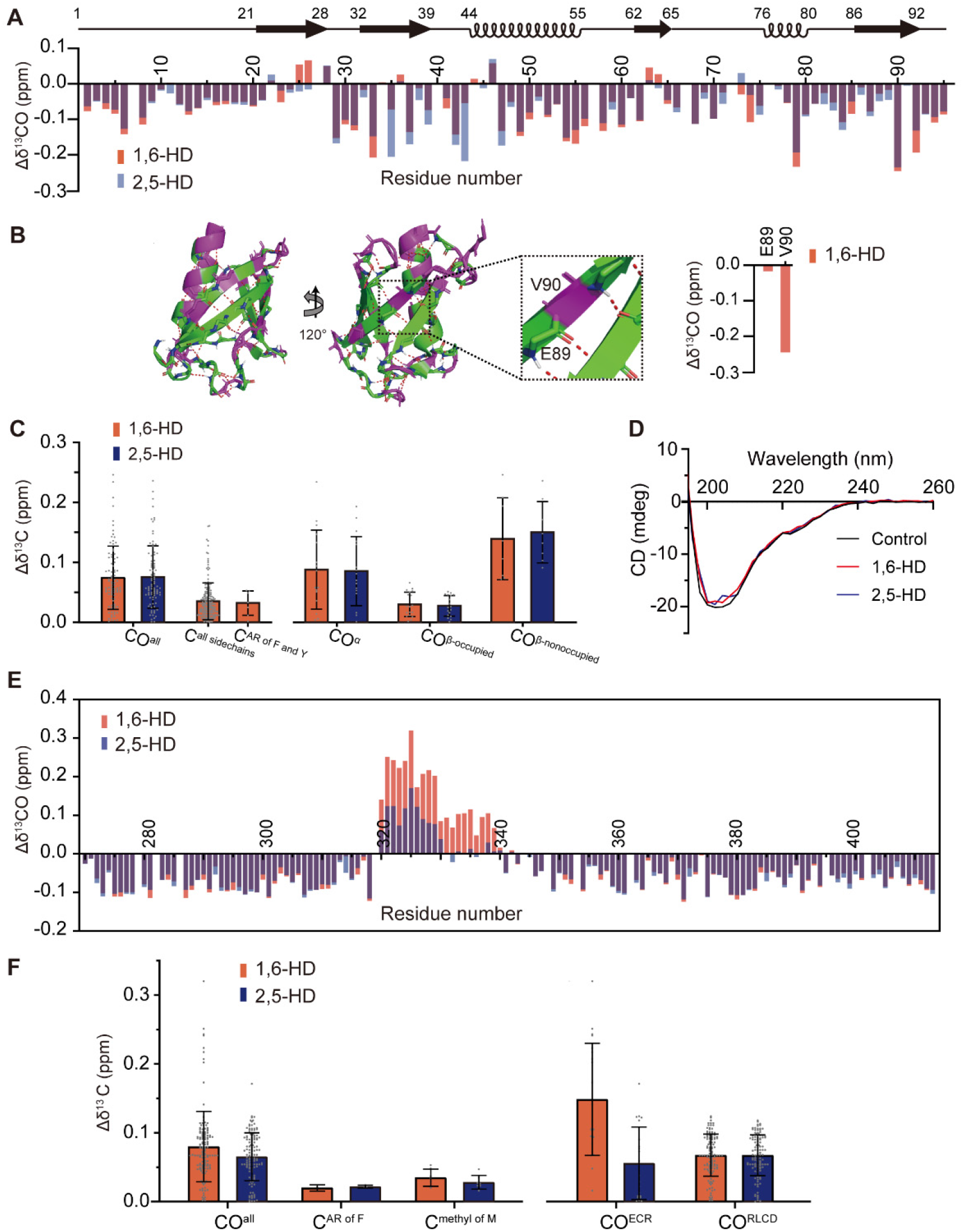
Solution NMR measurements of protein interaction by two aliphatic alcohols. Protein fragments corresponding to the well-folded SUMO1 protein (panels A-D) and the low complexity domain (LCD) of TDP-43 (panels E and F) were labeled with ^13^C and ^15^N, purified, incubated with or without 10% levels of either 1,6-HD (red) or 2,5-HD (blue), and visualized by solution NMR spectroscopy. (*A*) Chemical shift perturbations of carbonyl carbon atoms of the SUMO1 protein backbone in response to 1,6-HD (red) or 2,5-HD (blue) below schematic diagram of protein secondary structure (black). (*B*) Ribbon diagrams of SUMO1 protein structure displaying protein regions either highly perturbed (|Δδ^13^CO| > 0.07, purple) or not (|Δδ^13^CO| < 0.07, green) upon exposure to 1,6-HD. Expanded region shows the β-strand containing glutamic acid residue 89 (E89) and valine residue 90 (V90). The peptide backbone carbonyl oxygen atom of E89 is hydrogen bonded as part of the SUMO1 structure, and is insensitive to the effects of 1,6-HD (right side of panel). The carbonyl oxygen atom of V90 is free and shows a substantial chemical shift perturbation in response to 1,6-HD. (*C*) The combined chemical shift perturbations of carbon atoms from SUMO1 protein upon exposure to either 1,6-HD (red) or 2,5-HD (blue) for all backbone carbonyl groups (CO^all^) in HNCO, all sidechains including both aromatic rings and aliphatic sidechains (C^all sidechains^) that can be recognized in constant time ^1^H-^13^C HSQC, all aromatic rings (AR) of phenylalanine and tyrosine (C^AR of F and Y^) in constant time ^1^H-^13^C HSQC, backbone carbonyls in α-helix structures (CO^α^), backbone carbonyl groups in which the oxygen atom is hydrogen bonded as part of the SUMO1 protein structure (CO^β−occupied^), and backbone carbonyls in which the oxygen atom is free (CO^β−nonoccupied^). (*D*) Circular dichroism spectra of SUMO1 protein in the absence (black) or presence of either 1,6-HD (red) or 2,5-HD (blue). (*E*) Chemical shift perturbations of peptide backbone carbonyl carbon atoms of the TDP-43 LCD in response to 1,6-HD (red) or 2,5-HD (blue). Both aliphatic alcohols caused equivalent chemical shift perturbations to most carbonyl carbon atoms other than those residing within the evolutionarily conserved region (ECR) spanning residues 320-339. Within this region 1,6-HD caused more substantial chemical shift perturbations of carbonyl carbon atoms than 2,5-HD. (*F*) The combined chemical shift perturbations of carbons from TDP-43 LCD averages for all backbone carbonyls (CO^all^), all aromatic rings of phenylalanine residues (C^AR of F^), all methyl groups of methionine (C^methyl of M^), carbonyl groups limited to the evolutionarily conserved region of residues 320-339 (CO^ECR^), and carbonyl groups located throughout the remainder of the low complexity domain (CO^RLCD^).

Starting with the folded SUMO1 protein, we observed only modest, 1,6-HD-mediated chemical shift perturbations for amino acid side chains. More pronounced and equivalent chemical shift perturbations of backbone carbonyl groups were observed for both 1,6-HD and 2,5-HD. We interpret these chemical shift changes to be reflective of hydrogen bond interaction between hydroxyl groups of the aliphatic alcohols and carbonyl oxygen atoms of the SUMO1 polypeptide backbone (Fig. 2 *A-C*). Given that application of the aliphatic alcohols caused no more than modest chemical shift perturbations of the overall NMR spectrum, it is clear that neither chemical caused melting or denaturation of SUMO1 (*SI Appendix* Fig. S2). Such observations were confirmed by circular dichroism spectroscopy (Fig. 2*D*).

Chemical shift perturbations for carbonyl groups resulting from binding of 1,6-HD or 2,5-HD to carbonyl oxygen atoms of the SUMO1 polypeptide backbone were of two categories. Strong chemical shift perturbations were observed for carbonyls within β-strands wherein the oxygen atom was not hydrogen bonded as part of the protein structure. Substantially weaker chemical shift perturbations were observed for β-strand carbonyl oxygen atoms sequestered in the network of backbone hydrogen bonding holding β-strands together (Fig. 2 *B* and *C*). Strong chemical shift perturbations for non-occupied carbonyls within β-strands caused by 1,6-HD were also observed in the well-folded C2A and C2B domains of synaptotagmin1 (*SI Appendix* Fig. S3). Finally, clear evidence of both 1,6-HD and 2,5-HD binding was observed for many carbonyl oxygen atoms within both α-helical segments of the folded SUMO1 protein.

Turning to the LC domain of TDP-43, both aliphatic alcohols caused chemical shift changes for carbonyl groups of the polypeptide backbone, as well as more modest shifts of aromatic rings or methyl groups of amino acid side chains, including those from methionine residues (Fig. 2*E*, 2*F* and *SI Appendix* Fig. S4). The absolute values of these changes were larger for backbone carbonyl groups likely as a result of hydrogen bonding between hydroxyl groups of the aliphatic alcohols and carbonyl oxygen atoms of the polypeptide backbone. For most of the TDP-43 LC domain, 1,6-HD was no more active in this regard than 2,5-HD, nor was any appreciable difference observed between the regioisomers in binding the aromatic ring of phenylalanine residues or the side chains of other amino acids including methionine (Fig. 2*F* and *SI Appendix* Fig. S4).

Whereas negligible differences were observed in comparing the abilities of the two aliphatic alcohols to interact with backbone carbonyl oxygen atoms throughout most of the TDP-43 LC domain, isomer-specific differences were observed throughout the region spanning residues 320-339. Over the N-terminal half of this region, the 1,6-HD isomer elicited carbonyl group chemical shifts by roughly twice the magnitude of perturbation observed for 2,5-HD. Over the C-terminal half of this region, 1,6-HD effected changes in chemical shifts of backbone carbonyls, but 2,5-HD did not. It is precisely this region of the protein that is responsible for cross-β mediated self-association (10). This is likewise the region that adopts α-helical structure if the TDP-43 LC domain is studied at acidic pH in the absence of salt ions (11-13).

The Δ(*δ*Cα-*δ*Cβ) conformational changes revealed by comparison of chemical shifts observed in the presence of aliphatic alcohols with those of a random coil provide a measure of α-helical content (*SI Appendix* Fig. S5*A*). Positive Δ(*δ*Cα-*δ*Cβ) values are characteristic of α-helical conformation, whereas β-structures yield negative values (25). Hence, the observed Δ(*δ*Cα-*δ*Cβ) differences give indication that both 1,6-HD and 2,5-HD increase the population of α-helical structure in this region, and that the increase in α-helical conformation is slightly larger for 1,6-HD than 2,5-HD. Correspondingly, the CO conformational shifts calculated by comparison with random coil are also slightly more positive in the presence of 1,6-HD than of 2,5-HD (*SI Appendix* Fig. S5*B*). Hence, the different perturbations caused by the two aliphatic alcohols on CO chemical shifts (Fig. 2*E*) likely arise from a combination of effects due to both change in α-helical content as well as direct hydrogen bonding of diol hydroxyl groups to carbonyl oxygen atoms.

Considering that small effects may elicit more substantial consequences for cooperative processes including phase separation, these data can be interpreted to indicate that the stronger potency of 1,6-HD in disrupting TDP-43 self-association and phase separation, relative to 2,5-HD, may arise from either of two effects. Relative to 2,5-HD, the stronger-acting 1,6-HD regioisomer appears to prompt a slightly greater increases in α-helical content. Any condition favoring α-helical content in this region would of course impede cross-β interactions. 1,6-HD may likewise interact more effectively with backbone carbonyl groups than 2,5-HD within this localized region of the TDP-43 LC domain, thus preferentially inhibiting formation of the inter-strand hydrogen bond network required for cross-β interactions.

In summary, studies of aliphatic alcohol interaction with the folded SUMO1 and synaptotagmin1 proteins, as well as the LC domain of TDP-43, reveal pronounced chemical shift perturbations for carbonyl groups of the polypeptide backbone. We interpret these shifts to reflect interaction between hydrogen bond donors, in the form of hydroxyl groups of the chemical probes, and hydrogen bond acceptors in the form of carbonyl oxygen atoms of the polypeptide backbone. Significant differences in protein interaction by the two aliphatic alcohols was apparent in only one part of the TDP-43 LC domain, the ultra-conserved region of twenty residues known to mediate self-association and phase separation.

### Four Experimental Conditions Favor a β-to-α Conformational Switch

When incubated under conditions of physiologically normal salt and neutral pH, self-association of the TDP-43 LC domain leads to phase separation. Under these conditions, NMR signals broaden dramatically (Fig. 3*A*), hindering the use of triple resonance experiments to assign ^13^C chemical shifts that report on secondary structure. Three experimental approaches have mapped a cross-β forming region within the ECR of the TDP-43 LC domain. Two independent cryo-EM studies revealed virtually identical structures for the cross-β region between residues 314 and 327 (26, 27). This same region was mapped as the epicenter of the methionine oxidation-mediated footprint diagnostic of the TDP-43 domain whether assayed in hydrogel polymers, phase separated liquid-like droplets or living cells (9). Finally, the TDP-43 LC domain was probed via the systematic introduction of methyl caps upon nitrogen atoms. These methyl caps block the capacity for formation of individual, inter-strand hydrogen bonding of 23 consecutive residues within the ECR, thus yielding a modification-sensitive stretch of nine residues that mapped to the same location identified by cryo-EM structures and methionine oxidation footprinting (10).

**Fig. 3.**
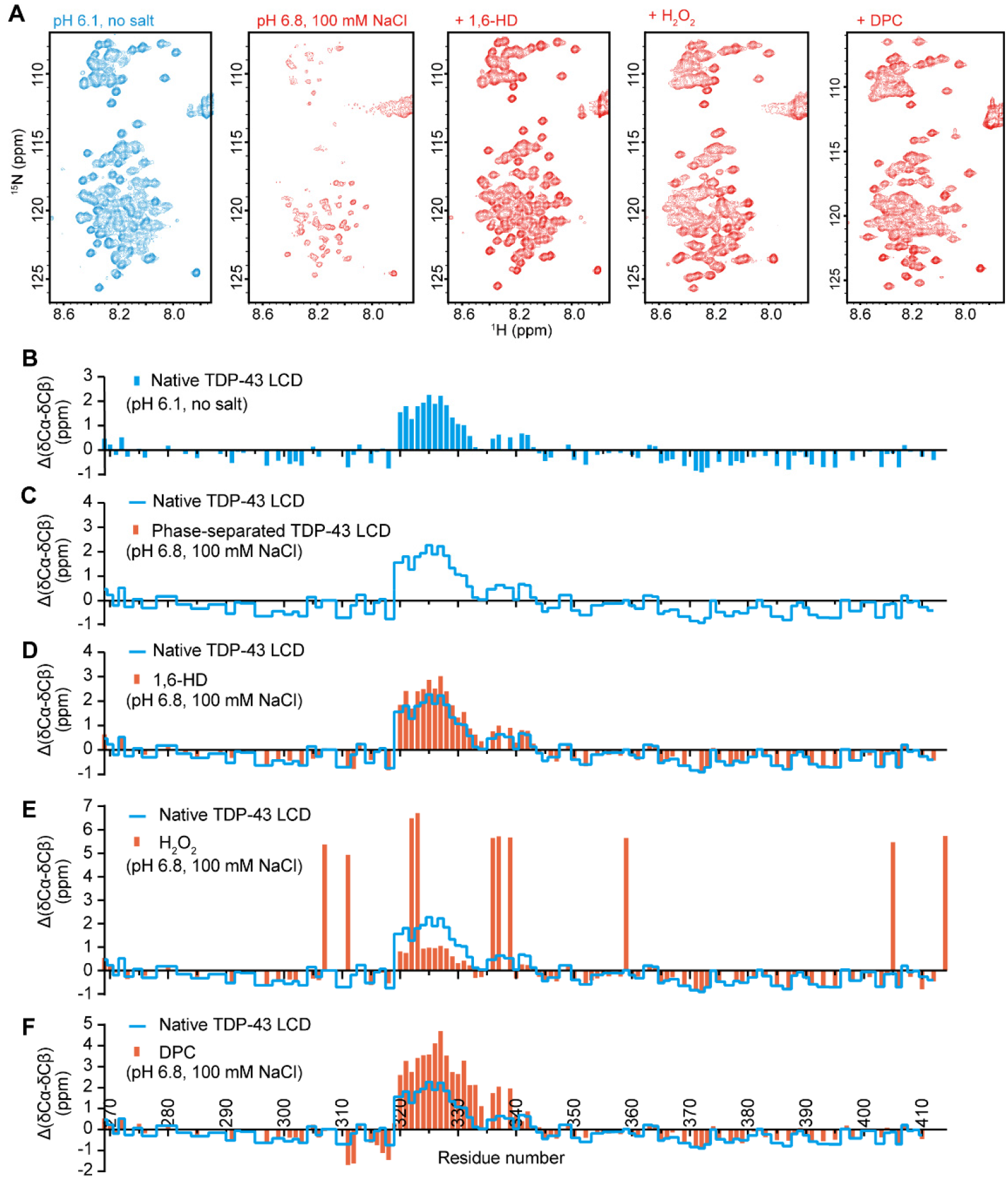
Solution NMR spectroscopic measurements of the TDP-43 low complexity domain under conditions that either allow or impede phase separation. (*A*) ^1^H-^15^N HSQC spectra of TDP-43 low complexity domain (LCD) under indicated conditions. Under conditions of acidic pH (6.1) and absence of salt ions, the TDP-43 LCD fails to undergo phase separation, thus allowing secondary structure analysis by solution NMR. Under conditions of pH 6.8 in the presence of 100 mM NaCl, the TDP-43 LCD undergoes phase separation, thus impeding NMR structural measurement, as indicated by the weak or absent ^1^H-^15^N HSQC cross-peaks. Upon exposure to 10% 1,6-HD, after H_2_O_2_-mediated methionine oxidation, or upon exposure to 80mM DPC, HSQC signals were restored, allowing NMR structural analysis. (*B-F*) Δ(*δ*Cα-*δ*Cβ) conformational shifts with respect to random coil of TDP-43 under different conditions: (*B*) Acidic pH in the absence of monovalent salt ions. Under such conditions the protein is unable to self-associate, thus allowing backbone resonance assignments with triple resonance experiments. The region between residues 320 and 339 is observed to adopt partially α-helical conformation as reported previously (11-13). (*C*) Backbone ^13^C chemical shifts cannot be obtained for phase separated protein as assayed in buffer conditions of neutral pH and physiologic levels of monovalent salt ions because triple resonance experiments are obscured by aggregation. (*D*) Protein at neutral pH and physiological levels of monovalent salt ions, yet exposed to 10% 1,6-HD. Evidence of α-helical secondary structure is observed between residues 320 and 339. (*E*) Protein at neutral pH and physiological levels of monovalent salt ions, yet exposed to 1% hydrogen peroxide (Materials and Methods). The large positive values of Δ(*δ*Cα-*δ*Cβ) arise because of oxidation of the ten methionine residues. Evidence of partial α-helical secondary structure is observed for the protein region between residues 320 and 330. (*F*) Protein at neutral pH and physiological levels of monovalent salt ions, yet exposed to 80mM the DPC lipid-mimic detergent (Materials and Methods). Evidence of increased α-helical secondary structure is observed between residues 320 and 339. For Panels C-F, blue lines correspond to a trace of the secondary shifts shown in Panel B.

If incubated in the absence of salt ions at an acidic pH, self-association and phase separation of the TDP-43 LC domain are fully prevented, thus allowing facile NMR analysis (Fig. 3 *A-C*). Under such conditions, as shown in Fig. 3*B*, the region of the protein between residues 320 and 339 exists in equilibrium between α-helical and random coil conformations. These data correspond well with earlier NMR studies of the TDP-43 LC domain conducted under conditions that prevent self-association and phase separation (11, 13).

Here we describe three additional conditions that inhibit TDP-43 LC domain self-association and concomitantly lead to the formation of α-helical conformation for the region between residues 320 and 339 (Fig. 3 and *SI Appendix* Fig. S6). The first means of effecting this β-to-α transformation in protein conformation was exposure to 1,6-HD. By inhibiting self-association and phase separation, exposure to 1,6-HD allows for solution NMR spectroscopy under physiological conditions of monovalent salt and neutral pH. Under such conditions, the region of the protein between residues 320 and 339 adopts a partially α-helical conformation characteristic of the protein when it is forced into the monomeric state at acidic pH in the absence of salt ions (Fig. 3*D*).

Methionine oxidation represents a second condition that prompts the β-to-α conformational switch. As described elsewhere, when phase separated preparations of TDP-43 LC domain are exposed to hydrogen peroxide, the liquid-like droplets melt in a manner correlated with methionine oxidation (9). This process is reversible. When the melted, oxidized sample is exposed to a mixture of two methionine sulfoxide reductase enzymes (28), thioredoxin, thioredoxin reductase and NADPH, the sample quickly re-forms phase separated droplets. Since methionine oxidation solubilizes the otherwise self-associated/phase separated TDP-43 LC domain, these conditions allow for solution NMR analysis. As shown in Fig. 3*E*, fully oxidized samples of the TDP-43 LC domain reveal formation of α-helical conformation in the ultra-conserved region.

The degree to which complete oxidation of all ten methionine residues within the TDP-43 LC domain prompts formation of α-helical secondary structure within the ultra-conserved region is reduced as compared with protein studied under conditions of acidic pH in the absence of salt ions, or under normal buffer conditions supplemented with 1,6-HD (Fig. 3 *B* and *D*). We hypothesize that this is reflective of oxidation of five methionine residues within the α-helical region (residues 322, 323, 336, 337 and 339). These methionine residues are substantially protected from oxidation if the protein is exposed to hydrogen peroxide in either its phase separated state or in living cells. Partial oxidation of the TDP-43 LC domain, limited primarily to methionine residues outside of the ultra-conserved region, effectively reverses self-association and phase separation (9).

To probe the idea that oxidation of methionine residues 322, 323, 336, 337 and 338 might inhibit formation of a helical secondary structure within the ultra-conserved region of the TDP-43 LC domain, we prepared protein samples bearing no oxidation, partial oxidation or complete oxidation of all ten methionine residues. ^1^H-^15^N HSQC spectra of each sample were acquired, allowing evaluation of the twenty residues localized to the α-helical region of the LC domain. As shown in *SI Appendix* Fig. S7*A*, complete oxidation of all ten methionine residues caused substantial shifts of the cross-peaks from the ultra-conserved region of the LC domain.

Partial oxidation led to a multiplicity of cross-peaks from this region which can be attributed to the existence of multiple species with distinct mixtures of both reduced and oxidized methionine residues (*SI Appendix* Fig. S7*B* and S7*C*). This cross-peak multiplicity hinders the assignment of ^13^C resonances to assess helicity, yet it is clear that the ^1^H-^15^N HSQC cross-peaks of the different, partially oxidized species are between the positions observed at pH 6.1/no salt and after full oxidation. As shown in *SI Appendix* Figure S7*C*, these cross-peaks are closer to the positions observed at pH 6.1/no salt. These observations can be explained by a model whereby there is equilibrium between random coil and α-helical conformations under each set of conditions, and that extensive oxidation decreases the population of the latter. The fact that the cross-peaks observed upon partial oxidation are close to those observed at pH 6.1/no salt shows that there is comparably robust formation of α-helical structure at partial oxidation levels fully adequate to impede phase separation (*SI Appendix* Fig. S7*B-E*).

The α-helix prone region of the TDP-43 LC domain contains seven alanine residues, five methionines and two leucines. As such, the exterior surface of the α-helix is hydrophobic. This hydrophobicity has been shown to favor binding of the lipid-like detergent dodecylphosphocholine (DPC). In a particularly thorough set of solution NMR experiments, it has been conclusively shown that exposure to DPC strongly enhances the proportion of the protein existing in the α-helical conformation (13).

Recognizing that lipid mimics favor α-helical conformation for residues 320-339, we tested the effects of DPC on self-association and phase separation of the TDP-43 LC domain. As shown in Fig. 4*A*, DPC dissolves phase separated droplets formed from self-associated TDP-43 LC domain. The droplet-melting effects of the lipid mimic are unique to the LC domain of TDP-43. Treatment of phase separated, liquid-like droplets formed from the LC domain of hnRNPA2 with DPC does not result in droplet melting (Fig. 4*B*).

**Fig. 4.**
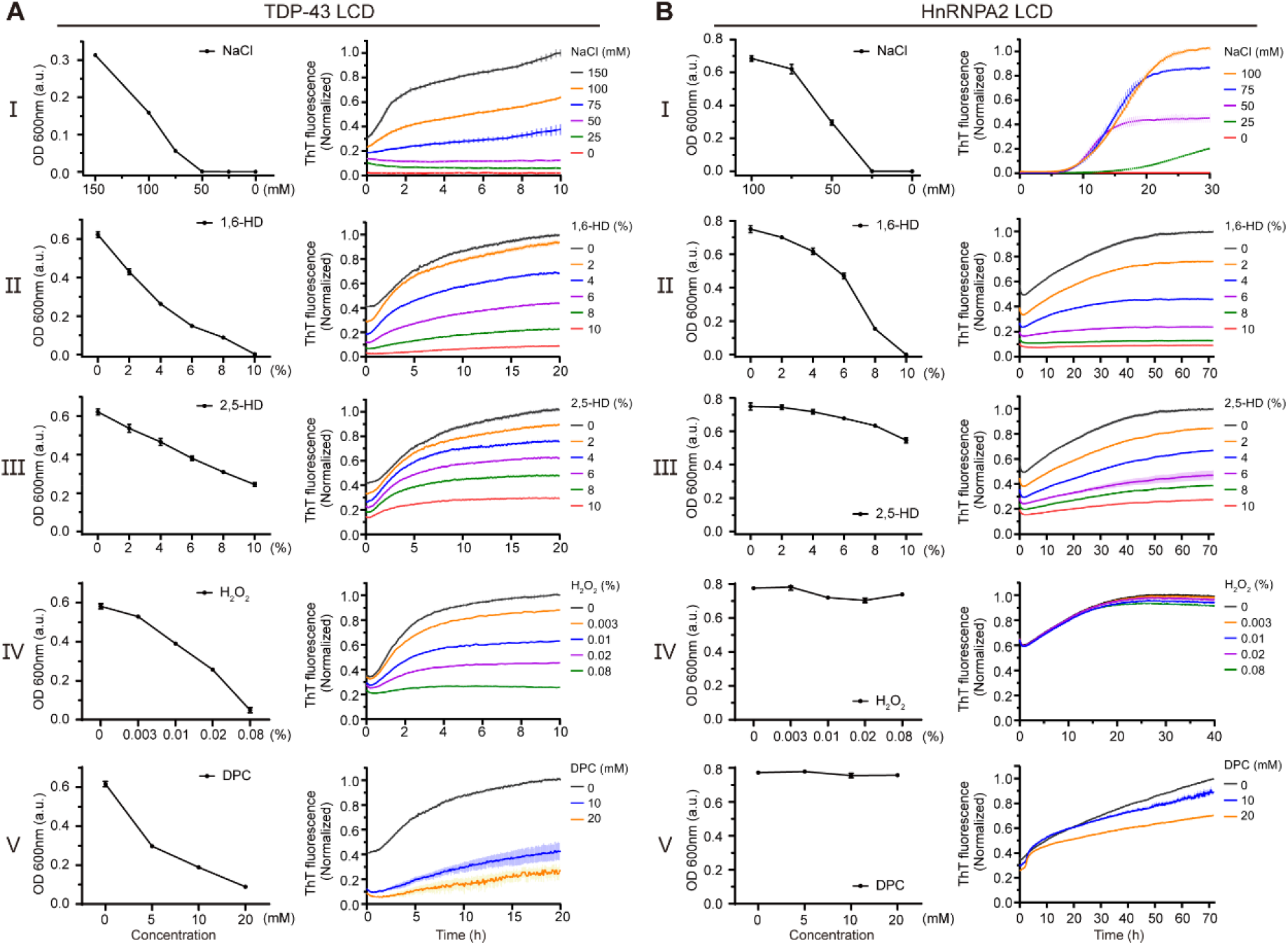
Effects of hypotonic buffer conditions, aliphatic alcohols, methionine oxidation and a lipid mimic on phase separation and cross-β oligomerization of the TDP-43 and hnRNPA2 low complexity domains. (*A*) A purified sample of the TDP-43 LCD was allowed to become phase separated in buffer of neutral pH and physiologically normal levels of monovalent salt prior to five conditions of experimental manipulation. Sequential reduction in monovalent salt concentration melted phase separated droplets as measured by turbidity (I). Exposure to 1,6-HD melted liquid-like droplets as a function of alcohol concentration (II). Diminished droplet melting was observed upon exposure to 2,5-HD (III). Significant droplet melting was observed upon exposure to hydrogen peroxide (IV) or non-ionic lipids (V). Formation of cross-β polymers by the same protein sample was monitored by acquisition of thioflavin-T fluorescence (Materials and Methods). Right Panels show impediments to cross-β polymerization effected by sequential reduction in monovalent salt concentration, exposure to 1,6-HD, exposure to 2,5-HD, exposure to hydrogen peroxide, or exposure to the lipid mimic DPC. (*B*) Parallel experiments performed with purified sample of the hnRNPA2 LCD. Reduction in salt concentration and exposure to aliphatic alcohols effected similar changes upon both phase separation and cross-β polymerization for the hnRNPA2 LCD as observed for the TDP-43 LCD. Neither hydrogen peroxide nor the DPC lipid mimic affected either phase separation or cross-β polymerization of the hnRNPA2 LC domain.

The ability of DPC to melt phase separated droplets of the TDP-43 LC domain formed under physiologic conditions of monovalent salt and neutral pH allowed samples to be evaluated by solution NMR spectroscopy. As shown in Fig. 3*F*, the resulting NMR spectra give clear evidence of α-helical structure in the protein region between residues 320 and 339.

### α-Helix Favoring Conditions Inhibit both Phase Separation and the Cross-β Conformational State

Having found four experimental conditions that allow formation of α-helical structure in the region of the TDP-43 LC domain spanning residues 320-339, we further asked what effects these conditions might have on the formation of cross-β structures. To this end we measured both phase separation and the acquisition of thioflavin-T fluorescence under five experimental conditions. The ground state condition employed normal levels of monovalent salt and neutral pH. To this we performed four experimental manipulations. First, we reduced ionic strength and shifted to buffer conditions of acidic pH. Second, we added sequentially increasing amounts of either 1,6-HD or 2,5-HD. Third, we added sequentially increasing amounts of hydrogen peroxide to effect methionine oxidation. And fourth, we added sequentially increasing amounts of the DPC lipid mimic.

As shown in Fig. 4*A*, all four of these manipulations impeded both phase separation, as monitored by the formation of liquid-like droplets, and binding of the thioflavin-T dye that favors interaction with cross-β assemblies (29). Of note, in comparing the two aliphatic alcohols, we observed that 1,6-HD impeded both assays at lower concentrations than 2,5-HD. We offer that these differences arise because of differential effects of the two aliphatic alcohols on the region between residues 320 and 339 of the TDP-43 LC domain, which may be reflected by different perturbations of carbonyl carbon chemical shifts of this region (Fig. 2*E*). It is precisely this region that forms strand-adhering cross-β interactions as deduced by structural biological studies (26, 27), protein footprinting studies performed both *in vitro* and *in vivo* (9), and biochemical studies using semi-synthetic proteins crafted to systematically mask single peptide nitrogen atoms across the ultra-conserved region (10).

Parallel experiments were conducted using a purified sample of the hnRNPA2 LC domain. Conditions of low salt and acidic pH blocked both phase separation and acquisition of thioflavin-T fluorescence by the hnRNPA2 LC domain (Fig. 4*B*). It was likewise observed that 1,6-HD was more effective than 2,5-HD in impeding both phase separation and thioflavin-T binding. By contrast, neither hydrogen peroxide nor the DPC lipid mimic exerted any detrimental effect on the ability of the hnRNPA2 LC domain to either phase separate or bind thioflavin-T. As such, we conclude that the latter sensitivities are unique to the LC domain of TDP-43.

## Discussion

In considering the biological utility of an LC domain regulated by methionine oxidation, we are influenced most heavily by published studies of the yeast ataxin-2 protein (7, 8). The LC domain of the yeast ataxin-2 protein contains 24 conserved methionine residues. The ataxin-2 LC domain self-associates via the formation of labile cross-β interactions, and these interactions are melted upon methionine oxidation. The yeast ataxin-2 protein resides in a cloud-like structure surrounding mitochondria, the integrity of which is regulated in living cells by the metabolic state of mitochondria. Oxidation of the yeast ataxin-2 LC domain weakens protein self-association and thereby transmits a signal of mitochondrial origin to the target-of-rapamycin (TOR) complex which, in turn, modulates autophagy.

Given that numerous reports have revealed close association of TDP-43 with mitochondria in vertebrate cells (30-32), we hypothesize that regulation of the oxidation state of the TDP-43 LC domain might also be controlled by the metabolic state of mitochondria. It is also well-established that TDP-43 is present in cytoplasmic RNA granules (33-35), and that 1,6-HD can melt these dynamic, membraneless organelles (36). Our simplistic belief is that methionine oxidation weakens self-association of the TDP-43 LC domain, thus favoring disassembly of RNA granules within which the protein resides. Finally, extensive evidence has been published revealing the integral role of TDP-43 in the control of localized translation (37, 38). Having found that exposure of the TDP-43 LC domain to hydrogen peroxide reverses self-association in a manner analogous to the activity of 1,6-HD, we offer that methionine oxidation may represent a biologically valid means of regulating the disassembly of RNA granules for localized de-repression of translation.

The cytoplasmic environment of all cells contains high concentrations of reduced glutathione and other oxidation scavengers that act to stringently prevent or reverse methionine oxidation, including two dedicated methionine sulfoxide reductase enzymes (28). As such, we anticipate that a specialized regulatory system will likely be involved in the redox control of methionine oxidation-mediated regulation of the TDP-43 LC domain.

Why would evolution co-specify the capacity to form two different protein conformations into the same, compact region of only 20 amino acid residues? We have described four different conditions that disfavor formation of TDP-43’s labile cross-β structure. These include acidic buffer conditions deprived of salt ions, exposure to aliphatic alcohols, exposure to non-ionic lipid mimics, and methionine oxidation. All four of these conditions concomitantly lead to the formation of a hydrophobic α-helix in precisely the same region of the protein used to form the labile cross-β structure require for LC domain self-association. Quite obviously, the α and β conformational structures formed by this region are mutually exclusive.

Whereas we can be confident of the biochemistry reported herein, we are uncertain of the biological utility of this oxidation-regulated conformational switch. If oxidation indeed takes place in proximity to mitochondria, it is possible that the hydrophobic α-helix might allow TDP-43 to associate with the mitochondrial membrane. If so, membrane interaction might cause TDP-43-bound mRNAs to be tethered to this cellular address. Indeed, biochemical fractionation experiments have revealed binding of endogenous TDP-43 to the inner membrane of mitochondria in neuronal cells (30). Figure 5 offers a graphical rendering of how we envision methionine oxidation to simultaneously dissolve RNA granules and allow the TDP-43 LC domain and its bound mRNAs to adhere to mitochondria.

**Fig. 5.**
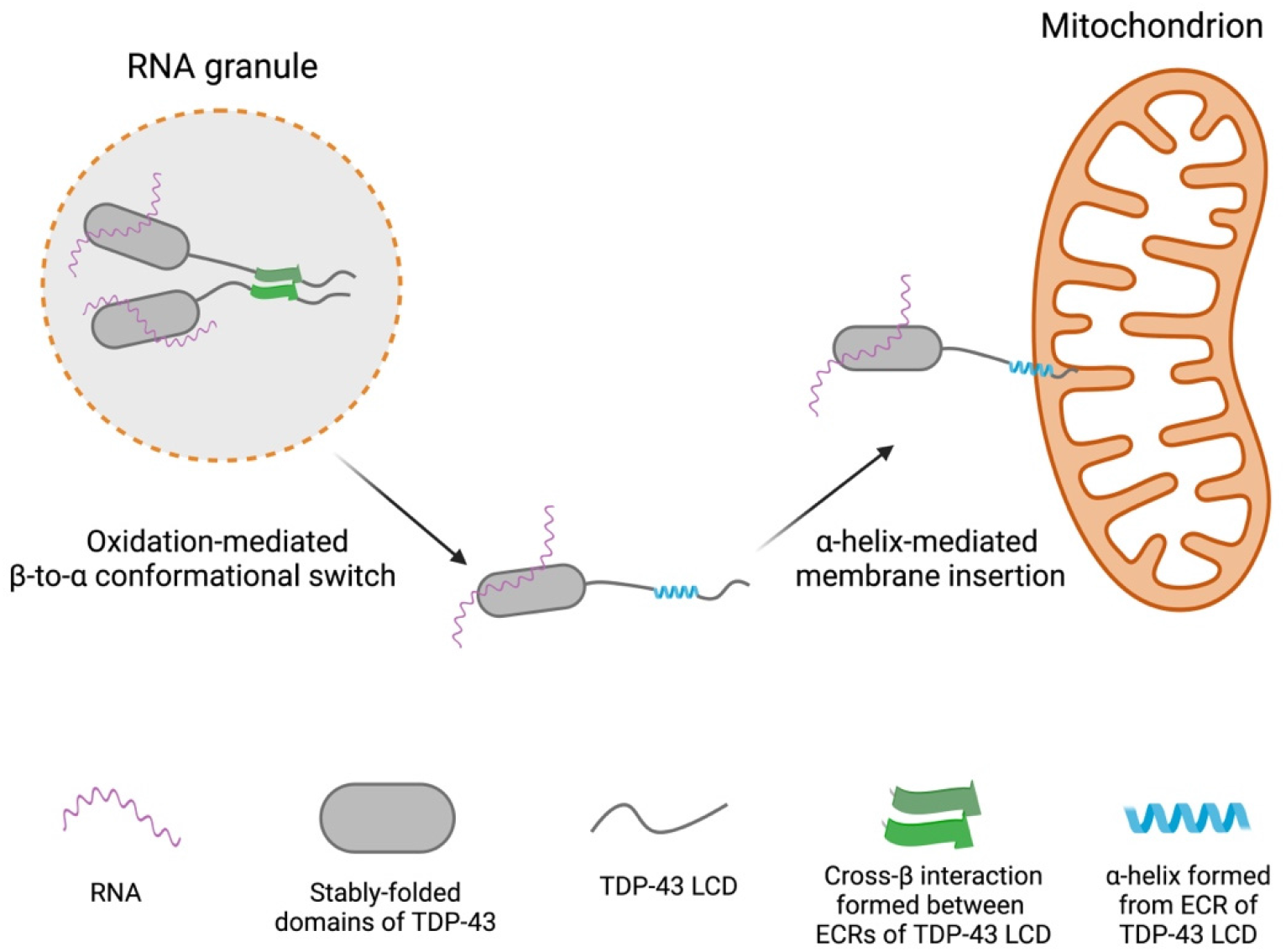
Conceptual diagram of methionine oxidation-mediated disassembly of RNA granules and adherence of released TDP-43/mRNA complex with mitochondria. Left image shows RNA granule containing TDP-43 in its self-associated state. Oxidation of methionine residues within the TDP-43 LC domain is hypothesized to dissolve the cross-β structure, allowing the evolutionarily conserved region (ECR) to assume an α-helical conformation. The extreme hydrophobic properties of the α-helix are then hypothesized to allow interaction with the membrane of mitochondria, thus resulting in local tethering of both TDP-43 and associated mRNA (right image).

If valid, these speculative ideas raise clear-cut questions. What interactions bring TDP-43-associated RNA granules into mitochondrial proximity, is there a TDP-43 receptor on the surface of mitochondria? What is the source of oxidation potential causing methionine oxidation? Once the hydrophobic α-helix of the TDP-43 LC domains is formed, is there an insertase machine required to embed it in the mitochondrial membrane (39)? Do the methionine sulfoxide reductase enzymes, in concert with thioredoxin, thioredoxin reductase and NADPH, reverse this oxidative switch and facilitate return of reduced TDP-43 to the nuclear compartment of cells?

We close with consideration of the experimental conditions hereby shown to disfavor formation of the labile cross-β structure responsible for self-association of the TDP-43 LC domain. Most importantly, because of its potential for biological relevance, how does methionine oxidation impede self-association? As for the aromatic phenylalanine and tyrosine residues known to be important for self-association of the FG repeats of nuclear pore proteins and the LC domains of many RNA-binding proteins (2, 5), we imagine that methionine residues help facilitate transient intermolecular contact between two separate LC domains. The weakly adhesive forces of methionine, tyrosine or phenylalanine residues are hypothesized to facilitate non-specific collision between LC domains residing on distinct polypeptide chains. Intermolecular collision events mediated by these weak and non-specific forces are required, we propose, for the formation of molecularly specific cross-β structures. Oxidation, we further propose, attenuates the hydrophobic character of the methionine side chain, thus disfavoring intermolecular collision of TDP-43 LC domains. Partial oxidation sufficient to fully disrupt phase separation does not destabilize the α-helical structure adopted by the ultra-conserved sequence as judged by the similarity of the ^1^H-^15^N HSQC spectra with that of the monomeric TDP-43 LC domain prepared in acidic buffer in the absence of salt ions (*SI Appendix* Fig. S7 *B* and *C*). Hence, such oxidation does not hinder the intramolecular forces required for α-helix formation despite eliminating cross-β mediated self-association and phase separation.

We offer similar thoughts as to why the DPC lipid mimic reverses the self-associated, phase separated state. Since the ultra-conserved region of the TDP-43 LC domain cannot simultaneously exist in both α and β conformations, helix-favoring lipids can be understood to drive equilibrium away from the labile cross-β structure required for self-association and phase separation.

Finally, the experiments reported herein offer a molecular explanation as to how two, regioisomeric aliphatic alcohols differentially melt disparate cellular structures reliant upon the self-adhesive properties of LC domains. As taught to us by Corey and Pauling over 70 years ago (40), it is the hydrogen bonding network mediated by peptide nitrogen atoms and carbonyl oxygens of the polypeptide backbone that are of paramount importance to the assembly of cross-β secondary structure. Elsewhere we have confirmed the importance of individual peptide nitrogen atoms along the polypeptide backbone of the ultra-conserved region of the TDP-43 LC domain for its ability to assemble into a labile cross-β conformation (10). As deduced by solution NMR spectroscopy, we now demonstrate that one of two related chemicals, 1,6-HD, perturbs carbonyl oxygens of the polypeptide backbone more readily than its regioisomer (2,5-HD). That differences in the abilities of the two regioisomeric chemicals to perturb carbonyl groups map precisely to the region that specifies self-association (residues 320-339) provides evidence that these differences represent the cause for the stronger relative activity of 1,6-HD in disrupting self-association and phase separation.

How do the regioisomeric chemicals differ in their effects upon self-association of the TDP-43 LC domain? On one hand, 1,6-HD increases α-helical content of the ultra-conserved region slightly more than 2,5-HD. It is possible that the tendency of this region to adopt an α-helical conformation emerged as a mechanism to limit the extent of cross-β structure formed by this region, thus yielding a dynamic apparatus relatively immune to aggregation. The slightly higher stabilization of α-helical structure caused by 1,6-HD compared to 2,5-HD may sufficiently tilt the equilibrium towards α-helical structure to disrupt phase separation owing to the cooperative nature of the process.

We also make note that the geometry of 1,6-HD, but not 2,5-HD, may allow for concomitant hydrogen bonding to two consecutive carbonyl oxygen atoms if the polypeptide backbone is disposed in an extended, β-strand conformation (16). Thus, 1,6-HD may interfere more effectively than 2,5-HD in blocking intermolecular interactions involving low populations of extended conformers that seed the formation of cross-β structure.

The solution NMR studies described herein provide evidence that, relative to amino acid side chains, 1,6-HD interacts preferentially with carbonyl oxygen atoms of the polypeptide backbone in both folded proteins and the disordered LC domain of TDP-43 (Fig. 2). Why, then, does 1,6-HD melt self-associated, phase separated LC domains while leaving folded proteins intact? We hypothesize that this stark difference owes to the fact that, in the case of LC domain self-association, aliphatic alcohols interfere with intermolecular interactions.

Once disrupted, these intermolecular interactions do not easily fall back into place. In the setting of well-folded proteins, impediments attributable to 1,6-HD interaction with the polypeptide backbone are limited to interference with intramolecular chemical bonds. Despite the widespread importance of backbone hydrogen bonding interactions to the stability of folded proteins, if transiently impeded by an aliphatic alcohol, we suggest that these intramolecular bonds quickly re-form.

Recent independent NMR studies focused upon the binding of 1,6-HD to the LC domain of the fused-in-sarcoma (FUS) RNA-binding protein likewise emphasize the importance of interaction of this aliphatic alcohol with carbonyl oxygen atoms of the polypeptide backbone (20). These concordant observations favor prediction that many cellular structures preferentially melted by 1,6-HD, relative to 2,5-HD, will be weakly adhered in a manner dependent upon the labile, cross-β self-association of LC domains. Knowing that the proteomes of eukaryotic cells contain thousands of LC domains, we predict that the weak interactions poised at the threshold of thermodynamic equilibrium described herein will be liberally deployed to assist in many forms of dynamic cell morphology and biologic function.

## Data, Materials, and Software Availability

All study data are included in the article and/or SI Appendix.

## Acknowledgements

We thank the Biomolecular NMR Facility at UT Southwestern Medical Center; Qiong Wu for technical help with NMR experiments; and Glen Liszczak and Deepak Nijhawan for scientific advice and encouragement. This work was supported by an anonymous donor (funds to S.L.M.); the National Institute of General Medical Science (grant GM130358 to S.L.M.); the National Cancer Institute (grant CA231649 to S.L.M.); and the Welch Foundation (grant I-1304 to J.R.).

