## Supplemental Information for "Oxidative regulation of TDP-43 self-association by a β-to-α conformational switch"

[1] Department of Biochemistry

[2] Department of Biophysics

UT Southwestern Medical Center

Dallas, Texas 75235

[3] Institute for Quantum Life Science

National Institutes for Quantum Science and Technology (QST)

4-9-1, Anagawa, Inage-ku, Chiba, JAPAN 263-8555

[#]authors contributing equally

### Supporting Information

#### Supplementary Figures

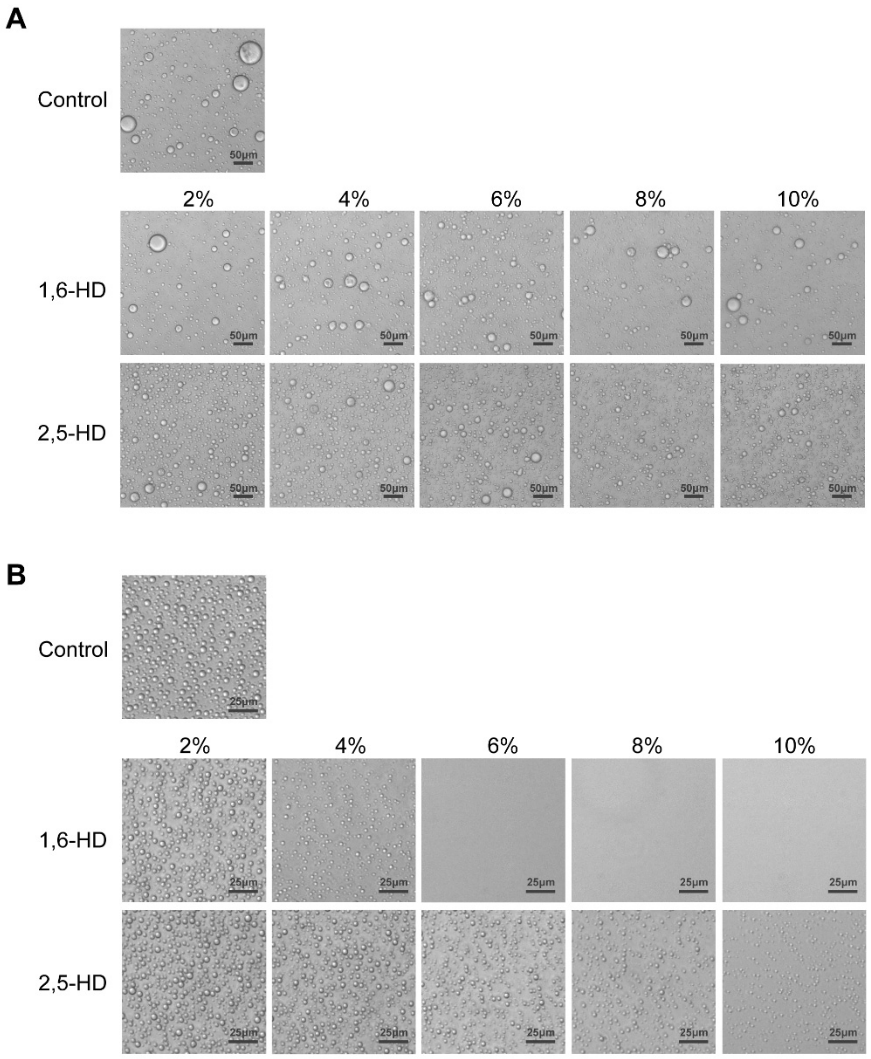


**Fig. S1. Effects of aliphatic alcohols on phase-separated liquid-like droplet formation by the structured and unstructured halves of TDP-43.**

**(A)** Liquid-like droplet formation by the structured N-terminal half of TDP-43 was induced by diluting the protein into the buffer in the presence 4% PEG-8000 and graded increases in concentration (w/v) of aliphatic alcohols as indicated. **(B)** Liquid-like droplet formation by the disordered C-terminal half of TDP-43 in buffer containing graded increases in concentration (w/v) of aliphatic alcohols as indicated. Scale bars = 25 µm.

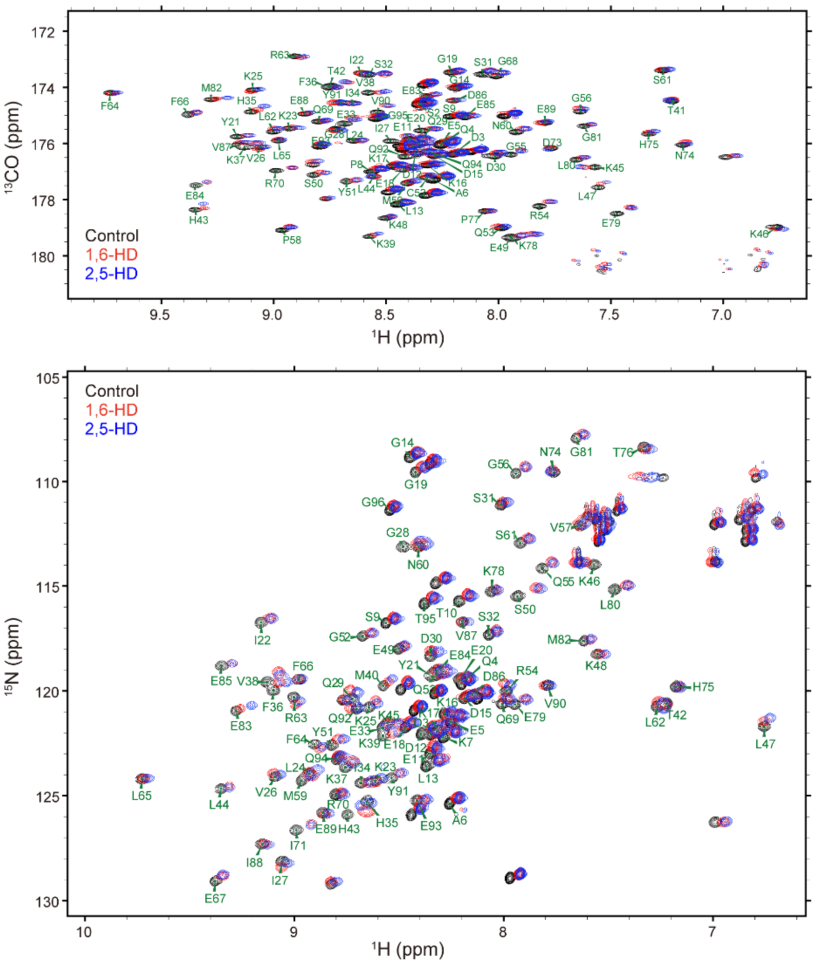


**Fig. S2. Chemical shift perturbation (CSP) analysis of the SUMO1 polypeptide backbone in the absence and presence of aliphatic alcohols.**

Diagrams show overlays of $^{1}$H-$^{13}$C HNCO spectra (upper panel) or $^{1}$H-$^{15}$N HSQC spectra (lower panel) of 150 μM SUMO1 obtained in NMR buffer alone (control, black), 10% (w/v) 1,6-HD (red), and 10% (w/v) 2,5-HD (blue). The NMR buffer consisted of 10 mM potassium phosphate, pH 6.5, 100 mM KCl, and 2 mM DTT (*1*).

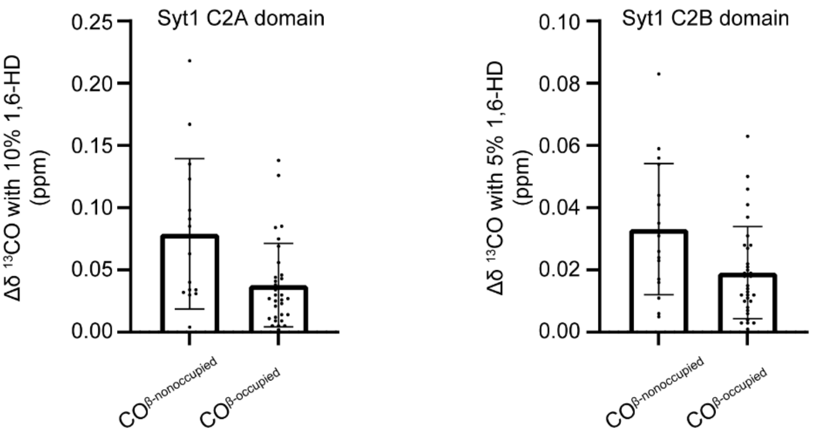


**Fig. S3. Effects of 1,6-HD on chemical shift perturbation of hydrogen bond-occupied or non-occupied carbonyl groups residing in β-sheets of the well-folded synaptotagmin1 C2A and C2B domains.**

Graphs show comparisons of absolute chemical shift perturbation (CSP) values of non-occupied and occupied peptide backbone hydrogen bonds as visualized by $^{13}CO$ in the presence of 1,6-HD. The CSP values are derived from all assigned residues from β-sheet structures of synaptotagmin1 C2A (left) and C2B (right) domains. Spectra were analyzed at 150 μM protein concentration in a buffer containing 50 mM HEPES, pH6.8, 100 mM NaCl, 2 mM DTT, and 1 mM EDTA (*2, 3*).

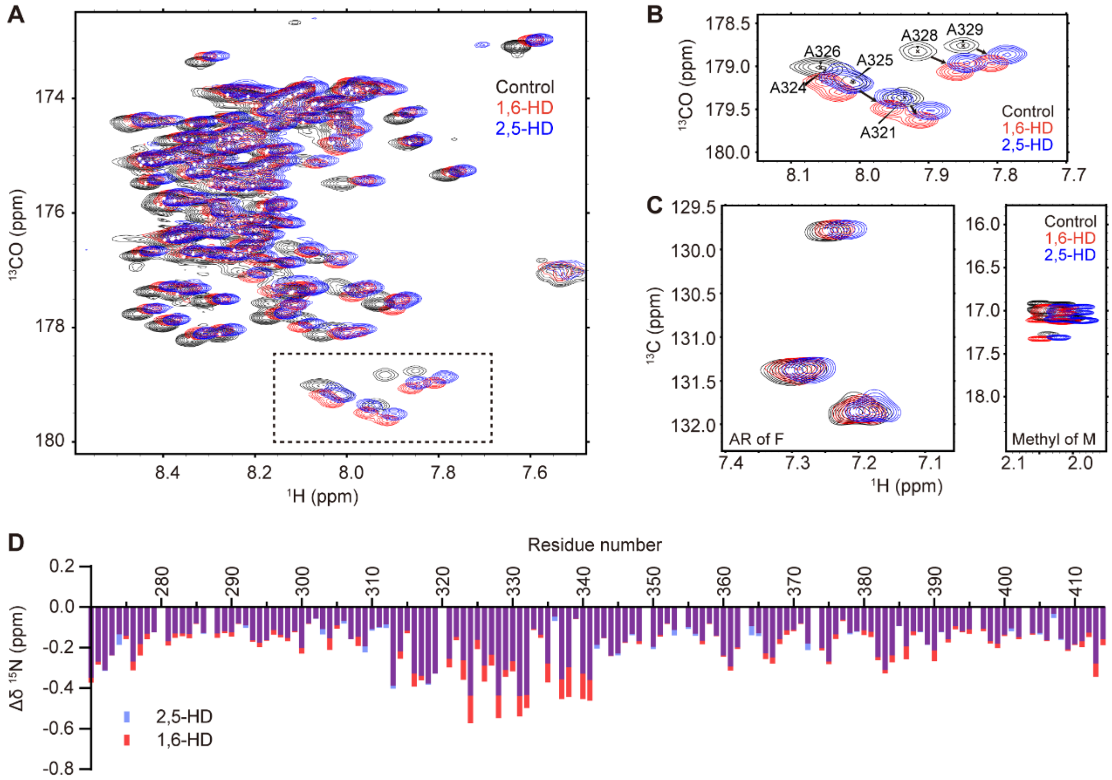


**Fig. S4. Chemical shift perturbation (CSP) analysis of TDP-43 low complexity domain (LCD) backbone and amino acid sidechains in the absence and presence of aliphatic alcohols.**

**(A)** Graph shows the overlay of 2D HNCO spectra of 100 μM $^{15}$N-/$^{13}$C-double-labelled TDP-43 LCD in a buffer containing 25 mM MES pH6.1 (black), or supplement with 10% (w/v) 1,6-HD (red) or 10% (w/v) 2,5-HD (blue). **(B)** Enlarged view of the area boxed with dash line in panel A. This area illustrates the CSP of Ala within TDP-43 LCD evolutionarily conserved domain. Black arrows indicate changes of cross-peaks of the indicated residues in the presence of aliphatic alcohols. **(C)** CSP of constant time $^{1}$H-$^{13}$C HSQC spectra of 100 μM Phe-[aromatic ring (AR), $^{13}$C6]-labelled TDP-43 LCD in the presence of 10% (w/v) aliphatic alcohols (left). The $^{1}$H-$^{13}$C HSQC spectra of 100 μM Met-[$^{13}CH_3$]-labelled TDP-43 LCD with/without 10% (w/v) aliphatic alcohols (right). NMR buffer: 25 mM MES pH6.1. **(D)** CSP of $^{15}$N atoms from the backbone of TDP-43 LCD with 10% (w/v) aliphatic alcohols at pH 6.1.

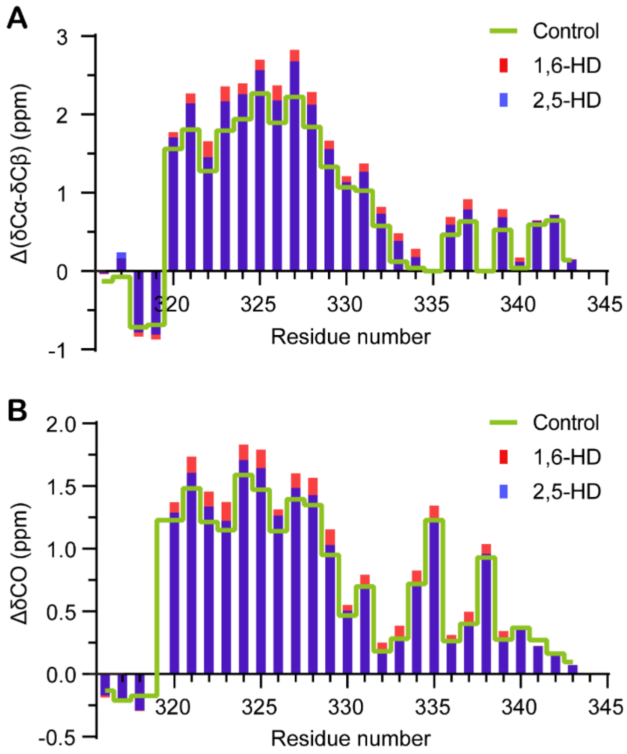


**Fig. S5. Aliphatic alcohols increase α-helical conformation of the evolutionarily conserved region (ECR) of the TDP-43 low complexity domain (LCD).**

**(A)** Secondary shifts (ΔδCα-ΔδCβ) of TDP-43 LCD (residue 316-343) in the absence or presence of 10% aliphatic alcohols in 25 mM MES buffer at pH6.1. The ECR refers to residues from 320 to 339. **(B)** Chemical shift deviations of backbone carbonyls of the TDP-43 LCD (residue 316-343) with respect to random coil. The ΔδCO represents the CO chemical shifts with or without 10% aliphatic alcohol treatment minus CO chemical shifts of the random coil in 25 mM MES buffer at pH6.1 (*4*).

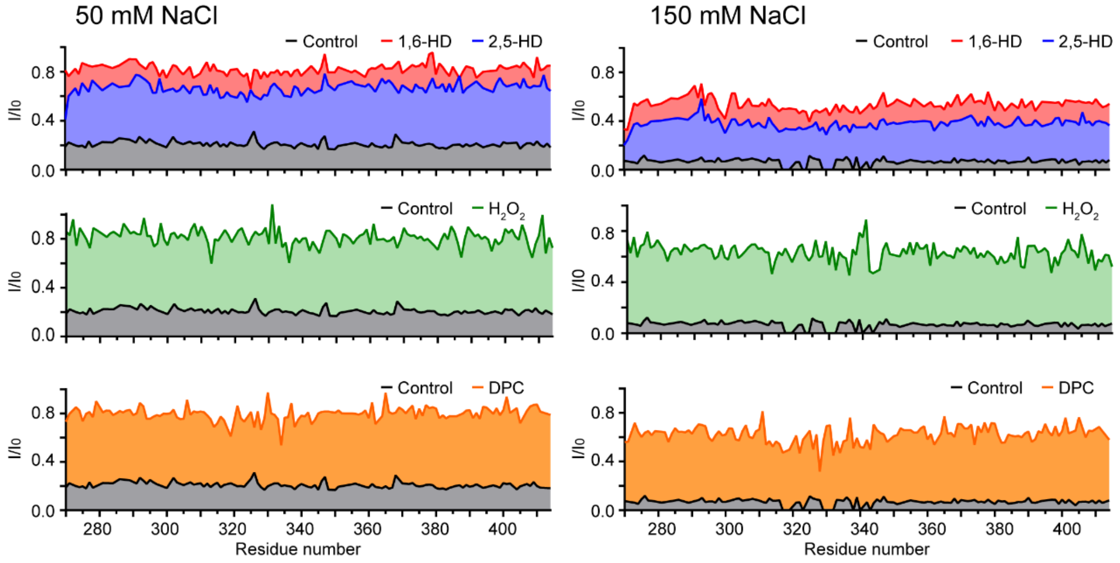


**Fig. S6. The effects of aliphatic alcohols, $H_2O_2$-mediated methionine oxidation, and exposure to the DPC lipid mimic on maintaining TDP-43 LCD $^1H$-$^{15}N$ HSQC signals in the presence of monovalent salt ions.**

$^1H$-$^{15}N$ HSQC signals (grey) of TDP-43 LCD drastically dropped due to self-association in buffer supplemented with 50 mM NaCl (left) or 150 mM NaCl-mediated phase separation (right). Exposure to 10% (w/v) of either 1,6-HD or 2,5-HD (top panel, red = 1,6-HD, blue = 2,5-HD) significantly enhanced NMR signals, as did $H_2O_2$-mediated methionine oxidation (middle panel), as did exposure to the lipid-mimic DPC (bottom panel). The $I_0$ represents the cross-peak intensity of native TDP-43 LCD, fully-oxidized TDP-43 LCD or native TDP-43 LCD with 200-fold DPC for each titration, under 40 μM $^{15}N$-labeled proteins, 25 mM MES (pH 5.5) without salts.

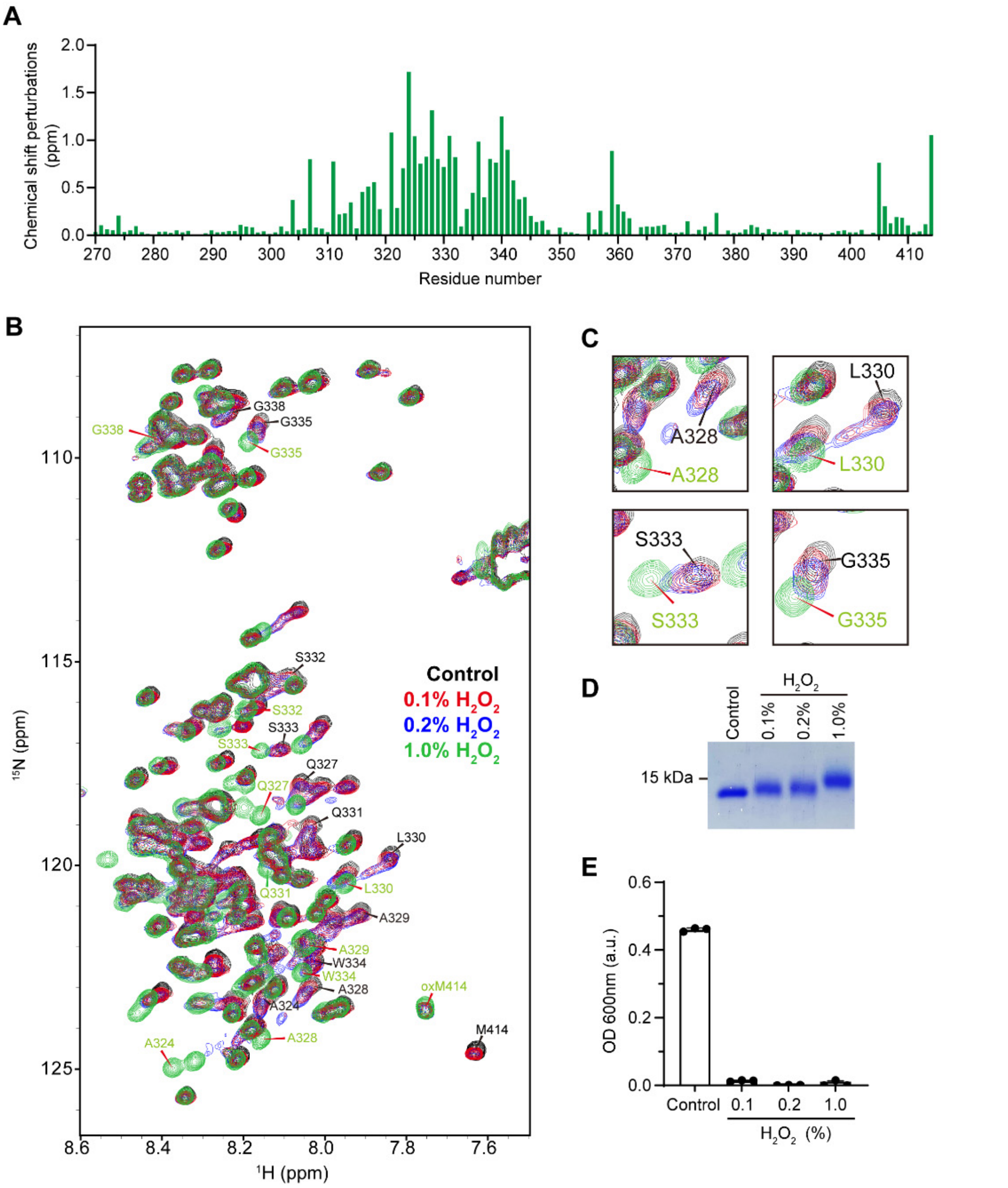


**Fig. S7. Partial oxidation retains α-helical structure of evolutionarily conserved region (ECR) of TDP-43 low complexity domain (LCD) while fully impeding phase separation.**

**(A)** The chemical shift perturbation (CSP) of fully-oxidized TDP-43 LCD on methionine (1% $H_2O_2$ for 30 min), compared to native TDP-43 LCD as assayed in buffer composed of 25 mM MES, pH6.1. The CSP was calculated by the following formula: $\sqrt{((\Delta\delta N)^2+(\Delta\delta HN)^2)}$. The graph shows significant CSP in TDP43-LCD ECR, representing the change of α-helix structures after full oxidation. **(B)** Overlay of $^{1}$H-$^{15}$N HSQC spectra of native TDP-43 LCD (control, black), partially-oxidized TDP-43 LCD (0.1% $H_2O_2$ red; 0.2% $H_2O_2$ blue), and fully-oxidized TDP-43 LCD (1% $H_2O_2$, green) in buffer composed of 25 mM MES, pH6.1. Representative residues are labelled in black (for control) or green (for 1% $H_2O_2$). For partially-oxidized samples, the majority of signals from the TDP-43 LCD ECR residues were retained near the signals of the native TDP-43 LCD, indicating that the partially-oxidized TDP-43 LCD retained higher population of α-helical structure compared with the fully-oxidized sample. The CSP of M414 represents an indicator of oxidation level. **(C)** Enlarged views show the CSP of A328, L330, S333, and G335 located within the ECR. **(D)** Coomassie stained SDS-PAGE showing the band shifts of TDP-43 LCD after partial and full oxidation. **(E)** Turbidity measurement of the samples in (B) at 50 μM protein concentration in buffer composed of 100 mM NaCl, 25 mM MES, pH 6.1. Fully reduced protein (control) was phase separated into liquid-like droplets. Exposure to 0.1%, 0.2% or 1% levels of $H_2O_2$ fully melted all liquid-like droplets.

#### Materials and Methods

##### Plasmid Construction

For *E.coli* expression, TDP-43 LCD (residue 263-414), and human SUMO1 (residue 2-97) were sub-cloned into pHis-paralle1 vector with a TEV cleavable site between protein and His tag. Fragments of TDP-43 NNR (residue 2-262), Mcherry-hnRNPA2 LCD (residue 190-341), MBP-hnRNPA2 LCD (residue 180-341), and hnRNPA2 LCD (residue 180-341) and were sub-cloned into pHis-paralle1 vector with caspase3 cleavable sites between protein and the His-fused tag.

##### Protein Expression and Purification

All the recombinant proteins were expressed in BL21 (DE3) competent cells, except expressing TDP-43 NRR in Rosetta (DE3) competent cells. The expression of TDP-43 LCD and hnRNPA2 LCD were induced with 1 mM IPTG at 37°C for 4 h. The expression of other proteins was induced by 0.5 mM IPTG at 18°C for 16-18 h.

Cell pellets expressing His-TDP-43 LCD were harvested and resuspended in lysis buffer (50 mM Tris, pH 7.5, 6 M guanidine-HCl (GdnHCl), 150 mM NaCl, 20 mM imidazole, 10 mM β-mercaptoethanol (BME)) and lysed by sonication for 3 minutes (10 seconds on/30 seconds off) on ice. The resulting cell lysate was centrifuged at 42,000 g for 50 min. The supernatant was loaded onto a gravity column with Ni-NTA resin (Gold Bio). The resin was washed by the lysis buffer, and then bounded protein was eluted with an elution buffer containing 50 mM Tris, pH 7.5, 6 M GdnHCl, 150 mM NaCl, 300 mM imidazole. The eluted protein was concentrated, flash frozen and stored at -80 degree. For NMR experiment, the eluted protein was further purified by HPLC and lyophilized. Protein powder was dissolved in pH3.5 water adjusted with 2N HCl and diluted into NMR buffer. To removes His tag from TDP-43 LCD, the water dissolved protein was diluted into 25 mM MES, pH 5.5 to a concentration of 15 μM and incubated with 30 μg/mL His-TEV enzyme at 37°C overnight. GdnHCl powder was added to the protein solution to a concentration of 6 M to completely dissolve and denature protein. The pH was then adjusted to 7.5 with 1M Tris (pH8.0) and the His-tag-cleaved TDP-43 LCD was clarified with Ni-NAT resin, concentrated, purified through HPLC, and lyophilized.

To prepare the fully-oxidized sample for NMR experiment, TDP-43 LCD was dissolved in 6 M GdnHCl buffer and incubated with 1% $H_2O_2$ for 30 min prior to HPLC purification.

The hnRNPA2 LCD was purified as described before (*5*). Briefly, the cells that expressed His-mCherry-caspase3-hnRNPA2 LCD were lysed in a buffer consisting of 50 mM Tris (pH 8.0), 2 M urea, 300 mM NaCl, 20 mM imidazole, and a Cooktail protease inhibitor by sonication (10 s on /1 min off) for 3 min in ice water. After centrifugation, the supernatant was loaded onto Ni-NAT resin. The protein was further eluted with a buffer consisting of 50 mM Tris (pH 8.0), 2 M urea, 300 mM NaCl, 300 mM imidazole. To remove the His-mCherry tag, the eluted protein was buffer exchanged into 50 mM Tris (pH 8.0), 2 M urea, 300 mM NaCl by 10 KDa concentrator, and further incubated with 3 μg/mL caspase3 enzyme for 3h at room temperature. The GdnHCl powder was added to the protein solution to a 6 M final concentration. Protein solution was subjected into Ni-NAT resin to clarify cleaved hnRNPA2 LCD. The resulting hnRNPA2 LCD was purified with HPLC, lyophilized, dissolved with pH 3.5 water or denaturing buffer, and then diluted into the buffer as indicated.

The SUMO1, TDP-43 NRR, and MBP-hnRNPA2 LCD were purified through similar methods. Cells were harvested and lysed in the buffer containing 50 mM Tris (pH 8.0), 500 mM NaCl, 20 mM imidazole, 10 mM BME, 5% glycerol, and Cooktail protease inhibitor by sonication (10 s on /1 min off) for 3 min on ice. For TDP-43 NRR, RNase was added to the lysis buffer. The supernatant was loaded onto Ni-NAT resin and unbound proteins were washed with the lysis buffer. The target proteins were eluted with 50 mM Tris (pH 8.0), 500 mM NaCl, 300 mM imidazole, 10 mM BME, 5% glycerol. The eluted SUMO1 protein was incubated with 6 μg/mL TEV and dialyzed against 50 mM Tris (pH 8.0), 200 mM NaCl, 2 mM BME overnight at 4 °C. The cleaved SUMO1 protein was then clarified with Ni-resin and further purified by size exclusion chromatography (SEC) (GE, Superdex 200pg 10/300 GL) in a buffer containing 12 mM phosphate buffer pH 6.5, 110 mM KCl, and 2 mM DTT. The eluted TDP-43 NRR was concentrated and further purified via SEC (GE, Superdex 200pg 10/300 GL) with a buffer containing 50 mM Tris (pH 8.0), 1 M NaCl, and 2 mM DTT. The MBP-hnRNPA2 LCD eluted from Ni-NAT resin was further purified by amylose resin. The final purified protein was concentrated and used in experiments immediately.

Synaptotagmin 1 C2A and C2B domains were purified as described before and stored in the NMR buffer containing 50 mM HEPES, pH6.8, 100 mM NaCl, 2 mM DTT and 1 mM EDTA (*2, 3*).

For universally labelled proteins used in NMR studies, they were expressed and grown in M9 minimal medium with $^{15}NH_4Cl$ (1 g/L) as the sole nitrogen source and [U-$^{13}$C6]-D-glucose (3 g/L) as the sole carbon source . Met-[$^{13}CH_3$]-labelled TDP-43 LCD was expressed in M9 minimal medium with 250 mg/L methyl-$^{13}$C-L methionine. Phe-[aromatic ring, $^{13}$C6]-labelled TDP-43 LCD was expressed in M9 minimal medium with all amino acids, including 200 mg/L [aromatic ring, $^{13}$C6]-Phenylalanine.

**Phase Separation Assay and Turbidity Measurement**

TDP-43 NRR-His phase separation/droplet formation was induced by diluting the protein to 20 μM concentration in a buffer containing 50 mM HEPES, pH 7.5, 100 mM NaCl, and 4% PEG 8,000 with absence or presence of indicated (hexanediols) HDs. Droplet formation of His-TDP-43 LCD was induced by diluting the protein to 15 μM concentration with a buffer containing 50 mM HEPES, pH 7.5, and 100 mM NaCl complemented with HDs. Images of liquid-like droplets were taken using Bio-Rad ZOE Fluorescent Cell Imager. Turbidity (OD 600nm) of the phase-separated sample was measured with a microplate reader.

To test effects of HDs and DPC on TDP-43 LCD and hnRNPA2 LCD phase separation, His-TDP-43 LCD and hnRNPA2 LCD were dissolved in 6 M GdnHCl buffer and diluted into a buffer containing 50 mM HEPES, pH 6,8 with presence of indicated HDs or DPC. The final protein concentration is 20 μM and the final GdnHCl concentration is 100 mM. To test effects of $H_2O_2$-mediated methionine oxidation on protein phase separation, the proteins were treated with 0-0,08% $H_2O_2$ as indicated in denaturing buffer for 45 min and diluted into the buffer containing 10 mM $Na_2SO_3$

used to quench $H_2O_2$. To test the effects of salt on protein phase separation, the lyophilized His-TDP-43 LCD and His-hnRNPA2 LCD powders were dissolved in water and diluted into the acidic buffer containing variant concentration of NaCl. Turbidity (OD 600nm) was measured to quantify phase separation.

**Thioflavin T (ThT) Fluorescence Kinetic Assay**

His-tagged TDP-43 LCD dissolved in denaturing buffer was diluted to 20 μM with a buffer containing 50 mM HEPES, pH 6,8, 50 μM ThT, 0.05% $NaN_3$, 100 mM GdnHCl, and HDs or DPC as indicated. MBP-hnRNPA2 LCD was used in ThT assay since it shows smaller variation than hnRNPA2 LCD alone under neutral pH condition. Fresh MBP-hnRNPA2 LCD stored in 500 mM NaCl was diluted to 50 μM with a buffer containing 50 mM HEPES, pH 6,8, 50 μM ThT, 0.05% $NaN_3$, 200 mM NaCl, and HDs or DPC as indicated. To prepare protein sample with $H_2O_2$-mediated methionine oxidation used in ThT assay, the proteins were partially oxidized by $H_2O_2$ for 45 min. $H_2O_2$ was then quenched with $Na_2SO_3$. To test effects of salt on cross-β polymer formation in ThT assay, His-tagged TDP-43 LC powder was dissolved in pH 3.5 water and diluted into an acidic buffer containing 20 μM proteins with variant NaCl concentration as indicated. HnRNPA2 LCD powder was dissolved into water and further diluted to 20 μM and complemented with variant amount of NaCl as indicated. All ThT samples were transferred to a 384- well plates (40 μL per well). The plate was sealed with a foil film to prevent evaporation. ThT fluorescence was monitored with excitation at 450 nm and emission at 485 nm at room temperature. The plate was shaken for 10 seconds prior to each measurement of ThT fluorescent intensity.

**Preparation of Partially Oxidized Samples**

In comparison to fully oxidized protein sample that was treated with 1% $H_2O_2$, the untagged TDP-43 LCD was partially oxidized by treating TDP-43 LCD polymers with less amount of $H_2O_2$. Briefly, the $^{15}N$-labeled TDP-43 LCD without His tag dissolved in pH 3.5 water was induced to undergo phase separation by diluting the protein to 200 μM with a buffer containing 50 mM Tris, pH 7.4 100 mM NaCl. The resulting phase-separated sample was then shaken overnight to product polymers at room temperature. The polymer was treated with 0.1% or 0.2% $H_2O_2$ for 30 min, and then mixed with $Na_2SO_3$ to quench $H_2O_2$. These partially oxidized polymers were collected by centrifugation (14000 rpm for 1h) and dissolved in trifluoroacetic acid (TFA). After blowing off TFA by air, the protein was dissolved in a denatured buffer containing 50 mM HAc-NaAc, pH 3.5 and 8 M Urea and dialyzed against pH 3.5 water overnight.

**Nuclear Magnetic Resonance (NMR)**

Backbone assignments of TDP-43 LCD, SUMO1, and Synaptotagmin1 C2A and C2B domains were accomplished according to methods reported in literatures (*1-3, 6*). The NMR experiments were performed at 25 °C on an Agilent DD2 spectrometer operating at 800MHz or 600MHz. The HNCO assay was used to determine the CSP (chemical shift perturbation) of $^{13}CO$ of $^{15}N$-/$^{13}C$-double labelled TDP-43 LCD and folded proteins under 10% (w/v) HDs. The constant time $^{1}H$-$^{13}C$ HSQC or $^{1}H$-$^{13}C$ HSQC assay was performed to detect the CSP of the protein sidechains. The same

NMR conditions with the assigned NMR spectra were used in the above assay (*1, 6*). NMR salt titrations for $^{1}H$-$^{15}N$ HSQC were performed under a condition of 40 μM $^{15}N$-labeled His-TDP-43 LCD, 25 mM MES (pH 5.5), 10% D2O, and presence of other indicated reagent. To determine the conformations of $H_2O_2$-oxidized TDP-43 LCD and native TDP-43 LCD with presence of 10% (w/v) HDs or 80 mM DPC, a pair of triple-resonance experiments HNCACB, CBCA(CO)NH were collected at indicated conditions for the sequential assignment of the $^{15}N$-/$^{13}C$-double labelled proteins (protein concentration is 400 μM). Untagged and $^{15}N$-/$^{13}C$-double labelled TDP-43 LCD was used in all triple-resonance experiments. All NMR data were processed by NMRpipe (*7*) and analysed by SPARK (*8*).
